# GenOO: A Modern Perl Framework for High Throughput Sequencing analysis

**DOI:** 10.1101/019265

**Authors:** Manolis Maragkakis, Panagiotis Alexiou, Zissimos Mourelatos

## Abstract

**Background:** High throughput sequencing (HTS) has become one of the primary experimental tools used to extract genomic information from biological samples. Bioinformatics tools are continuously being developed for the analysis of HTS data. Beyond some well-defined core analyses, such as quality control or genomic alignment, the consistent development of custom tools and the representation of sequencing data in organized computational structures and entities remains a challenging effort for bioinformaticians.

**Results:** In this work, we present GenOO [jee-noo], an open-source; object-oriented (OO) Perl framework specifically developed for the design and implementation of HTS analysis tools. GenOO models biological entities such as genes and transcripts as Perl objects, and includes relevant modules, attributes and methods that allow for the manipulation of high throughput sequencing data. GenOO integrates these elements in a simple and transparent way which allows for the creation of complex analysis pipelines minimizing the overhead for the researcher. GenOO has been designed with flexibility in mind, and has an easily extendable modular structure with minimal requirements for external tools and libraries. As an example of the framework’s capabilities and usability, we present a short and simple walkthrough of a custom use case in HTS analysis.

**Conclusions:** GenOO is a tool of high software quality which can be efficiently used for advanced HTS analyses. It has been used to develop several custom analysis tools, leading to a number of published works. Using GenOO as a core development module can greatly benefit users, by reducing the overhead and complexity of managing HTS data and biological entities at hand.

## BACKGROUND

The recent development of high-throughput sequencing (HTS) technologies has significantly lowered the cost of sequencing through parallelization, making HTS a widely used tool for genomic analysis. A typical HTS run involves the sequencing of millions of short reads that are usually parts of larger entities such as chromosomes or RNA molecules. There are several types of experiments involving HTS, such as Whole Transcriptome Shotgun Sequencing (RNA-Seq) [1], which has become the de-facto experiment for transcriptomics analysis. High-throughput sequencing of RNA isolated after CrossLinking and ImmunoPrecipitation [2] (HITS-CLIP, CLIP-Seq) and Photoactivatable-Ribonucleoside-Enhanced Crosslinking and Immunoprecipitation [3] (PAR-CLIP) are common techniques used to identify RNA binding sites for RNA binding proteins (RBPs). Similarly, ChIP-Seq [4] is a widespread technique in which genomic locations bound by DNA binding proteins, such as transcription factors and histones, are sequenced. The number of HTS methods is ever expanding, highlighting the need for efficient computational analysis of the datasets produced.

In a typical HTS experiment the sequenced reads initially undergo pre-processing steps in which adaptor sequences may be removed, low read quality nucleotides can be trimmed, and several quality control tests executed. Next, the reads are aligned to a reference genome or transcriptome, or can be used for genome/transcriptome assembly. Several tools with well-defined scope can handle specific parts of the sequencing analysis. For example, the alignment step can be performed by each of several programs such as BWA [5] or Bowtie [6]. Other tools such as Cufflinks [7] can be used to assemble transcripts and estimate their relative abundance based on RNA-Seq data.

While the core of a HTS analysis consists of these basic well defined steps, it is often the case that custom analyses need to be performed either alongside or following these steps. These custom analyses often directly address biologically relevant questions and tend to be highly specific and diverse. To date, several code libraries have been published attempting to facilitate the development of tools to be used in such analyses, such as BioPerl [8], BioPython [9] and BioRuby [10]. Due to the fact that these libraries have been designed for broad and generic use, they can be particularly complicated and intimidating for novice users, and additionally notoriously hard to manage for more advanced users and code contributors.

In this paper we present GenOO, an open-source Perl framework specifically designed for the development of HTS analyses. The framework is developed around Moose, a modern object system for Perl. Perl has been established as one of the leading programming languages in the bioinformatics community, as its features and string processing capabilities make it an ideal solution for developing bioinformatics applications. Though Perl has been criticized for its cumbersome object system in the past, the introduction of Moose has provided the language with a very concise and appealing object system that makes writing object oriented code effortless. The GenOO framework uses modern Perl concepts such as roles, to avoid common pitfalls such as deep inheritance trees. It models biological entities and core HTS structures into Perl objects. It also provides classes and methods to easily manipulate common file types used in sequencing analysis and supports the use of databases to tackle the enormous amounts of data produced by high throughput sequencing experiments. However, arguably the most important feature in GenOO is that all the above mentioned modules are integrated with each other enabling advanced queries and analysis that combine several of the modules simultaneously. In essence, GenOO provides a feature-rich, easy-to-use, consistent programming interface for custom HTS analysis. The primary focus during the framework design was to make it easily extensible and flexible without sacrificing optimization and performance.

In the following we also present a simple yet realistic example of the framework’s flexibility and capabilities by presenting a custom use case of HTS analysis and how this can be implemented using GenOO. Specifically, we present a small script that would filter a BED format file of Single Nucleotide Polymorphisms (SNPs) to keep only those that are contained within the coding sequence of expressed mRNAs. The expressed mRNAs will be defined in respect to their RPKM[11] values as defined by an RNA-Seq experiment. The RPKM measure is a simple calculation used to present the capabilities of the framework. In a real analysis different normalization techniques can easily be substituted.

## IMPLEMENTATION

### Basic Concepts and Structural Elements

The GenOO framework is developed around Moose, a widely used modern object system for Perl 5 that enhances the writing of object-oriented Perl. Moose greatly simplifies the way object oriented code is written in Perl and is particularly easy for novices to master, as it borrows many useful features from programming languages such as Perl 6, Lisp, Java, and Ruby. We have used Moose as the base for the implementation of all GenOO classes, allowing for concise, flexible and extensible code.

In contrast to Perl 5, Moose extensively uses roles as an alternative to deep hierarchies and base classes. A role encapsulates some piece of behavior or state that can be shared between classes. Roles are said to be “consumed” by other classes and all the methods and attributes of the role are available to be used by the class as if they have been defined in the class itself. Roles are similar but not equivalent to mixins and interfaces of other programming languages, as they can also require consuming classes to implement certain methods.

In GenOO we follow the dependency injection principle which is a software design pattern that removes hard-coded dependencies from within the classes and makes it possible to change them, either at run-time or at compile-time. Practically, following this principle we avoid using hard coded constructor calls to external classes within any class method. We design each class so that it specifically asks for all its requirements (instances of other classes) through its own constructor and does not rely on instantiating a hard coded one. Dependency injection greatly increases code modularity and testability, facilitating extension and code reuse. As a result of this design principle, object instantiation is mainly performed through factory classes following the factory design pattern.

The factory design pattern is one of the most commonly used design patterns in modern programming. Its aim is to create objects without exposing the instantiation logic to the client, but instead refer to the created object through a common interface. When a client needs a product, it initiates a request to the factory for a new product, providing the information about the type of the object it needs. The factory instantiates a new concrete product and returns it to the client. The client uses it as an abstract product without being aware of its concrete implementation. The advantage of using the factory design pattern is that it increases code flexibility, as new classes can be easily added, and existing classes modified, without changing the framework code which relies on them.

To support further development and improvement of the framework, we have implemented an extended test suite. The suite is based on object-oriented code and extensively covers the framework’s functionality, providing a safeguard for further refactoring and development.

### Design

Arguably, the backbone of the GenOO framework is the Region role which corresponds to a generic area along a reference sequence. The role requires consuming classes to implement certain attributes (i.e. “strand”, “rname”, “start”, “stop”, “copy_number”) and provides additional advanced methods such as the distance from another region for free. This role is consumed by multiple other classes within the framework, including the classes that correspond to HTS reads, and provides a common base for code integration (Figure 1) as it enables all modules to speak in a common language.

**Fig. 1.**
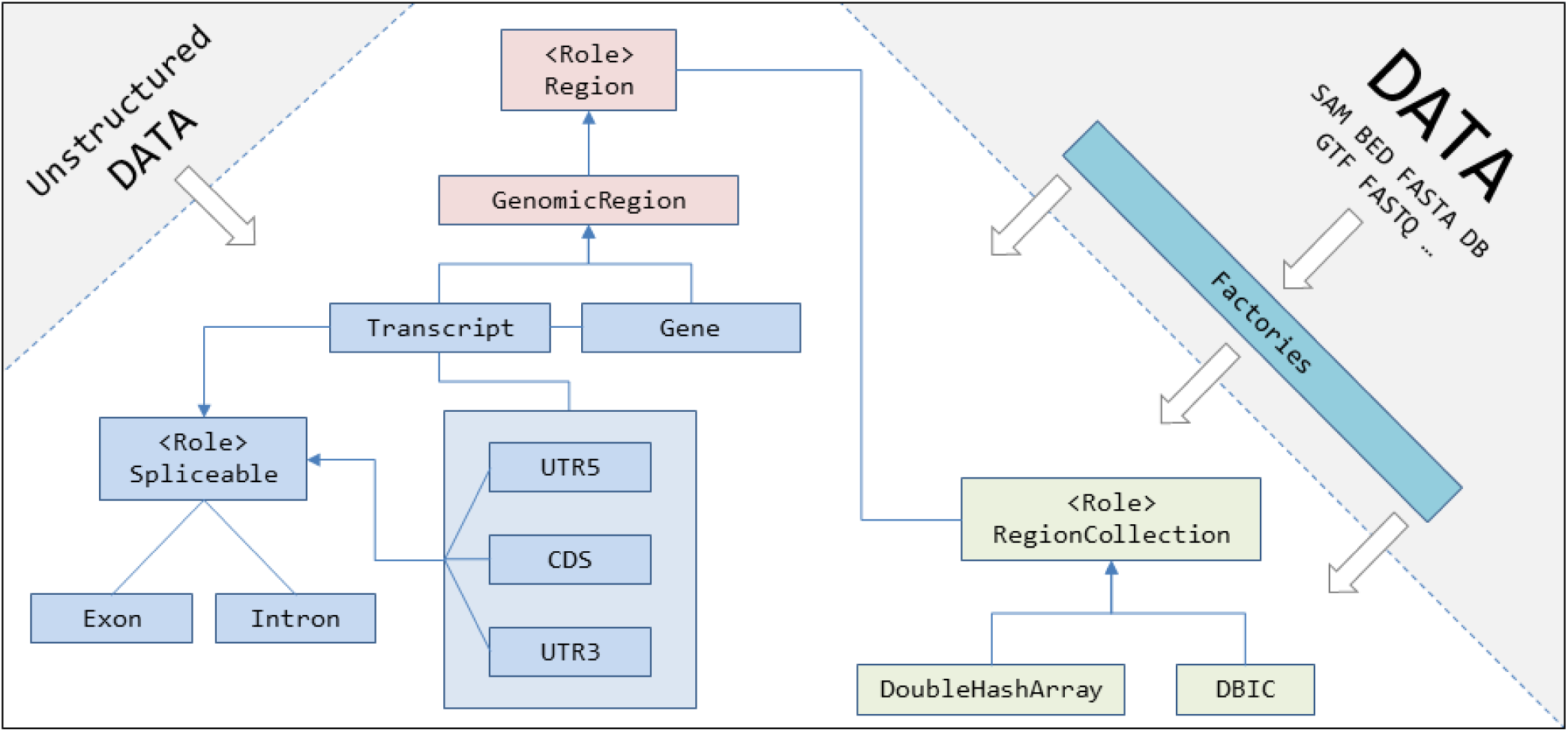
The basic class structure of GenOO. The input for structured data in known file formats is done through factory classes. Unstructured data can be introduced directly to class constructors. Arrows represent inheritance or role composition. For example the `GenomicRegion` class consumes the `Region` role. Solid line connectors represent association of classes. For example a transcript belongs to a gene and a gene may have multiple transcripts.

The GenomicRegion class consumes the generic Region role and specifies the reference sequence as a particular chromosome. GenomicRegion also has the “species” attribute which enables genomic analysis for different species simultaneously. The GenomicRegion serves as the base class for more advanced classes that correspond to specific genomic elements along the genome such as genes, gene transcripts and others (Figure 1).

The Transcript class corresponds to a gene transcript and usually belongs to a Gene class. The Gene class, in essence, is defined as a collection of Transcript objects. These two classes are designed so that they can communicate with each other in order to extract required information. Transcripts are divided into protein coding and noncoding ones, protein coding ones having methods that can extract the coding (CDS), 5’ UTR (UTR5) and 3’UTR (UTR3) sequences and coordinates (Figure 1).

A particularly important structure within the genomic group of classes is the Spliceable role. Spliceable provides functionality for entities/classes that undergo splicing. The most critical such class is the Transcript but this role is also composed into the UTR5, UTR3 and CDS classes. The use of the Spliceable role within these classes greatly reduces deep hierarchy and encapsulates the piece of behavior that is shared between them. Spliceable supports several advanced methods such as the extraction of exonic and intronic sequences and facilitates the management of the complex structure.

In HTS it is often the case that a user needs to perform operations on groups of elements and in particular regions. In GenOO we have implemented the RegionCollection role as an interface for classes that serve as a collection of regions. One of the most common analyses is querying for regions that fulfill certain positional criteria within a collection. Perhaps, the most characteristic such query is the selection of regions that overlap with another one. This query can be particularly demanding computationally, if implemented with a naive, brute force approach. In GenOO, we have implemented a collection engine that supports such queries and is explicitly named as DoubleHashArray after the data structure that it uses. This structure tackles this computational problem by storing data in a two dimensional Perl hash using “strand” as a primary key and “chromosome” as the secondary key. Each such key pair points to the sorted array of regions that are found in that particular strand and chromosome. The array is sorted by ascending “start” values of the included regions. When the query is performed the two keys are used to rapidly locate the array with all the regions on that specific “strand” and “chromosome”. This step in principle corresponds to partition pruning performed by database engines (e.g. MySQL). Following this step, a binary search algorithm is used to locate the regions of the array that satisfy the given positional criteria. This technique reduces computational complexity and allows for the rapid extraction of the result. Although DoubleHashArray can be quite fast, it suffers from the fact that the data need to be in memory, and therefore it can only be used for relatively small data sets, and mostly for prototyping and draft solutions. In contrast, we have also implemented a pure database oriented collection engine which is named DBIC. Currently, the implemented classes support database tables with the following columns: “strand”, “rname”, “start”, “stop”, “copy_numbef’, “sequence”, “cigar”, “mdz”, ”number_of_best_hits” but the user can easily extend them to support any table schema provided that it supports all columns/attributes defined in the Region role. The GenOO DBIC class is based on DBIx::Class which is a modern Perl module that provides an extensible and flexible object-relational mapper. DBIx::Class supports most major databases such as SQLite, MySQL, PostgreSQL and Oracle.

For Input/Output we have implemented several file parsers in pure Perl for the most common HTS file formats such as BED, FASTA, FASTQ, SAM and others. External requirements have been limited to the absolute bare minimum in order to facilitate code deployment and expandability. All file parsers are fully compatible and integrated with the rest of the GenOO framework and can easily fit into any complex analysis. For example, parsing a SAM formatted file line by line returns a SAM record which consumes the Region role and can be directly inserted into a RegionCollection instance or can be used to query a collection of genes or transcripts to efficiently get the elements that it overlaps with. The framework seamlessly supports both gzipped and plain text files. To enhance code simplicity and structure we have developed several factory classes that create simple and complex RegionCollection instances from the most common file types.

### Use Case

In figure 2 we present an example of the framework’s flexibility and capabilities by programming a custom but realistic use case of HTS analysis. We want to filter a list of Single Nucleotide Polymorphisms (SNPs) keeping only those that are contained within the coding sequence of expressed mRNAs that are expressed above a specific expression level, as defined by an RNA-Seq experiment. To keep the example simple we define expressed mRNAs as those with RPKM values greater than a user defined value. We also assume that through some initial pre-processing steps the RNA-Seq reads have been aligned to a reference genome and have been stored in a database table. Under different circumstances and using slight modifications the RNA-Seq reads could be easily be read from a SAM file or similar.

**Fig. 2.**
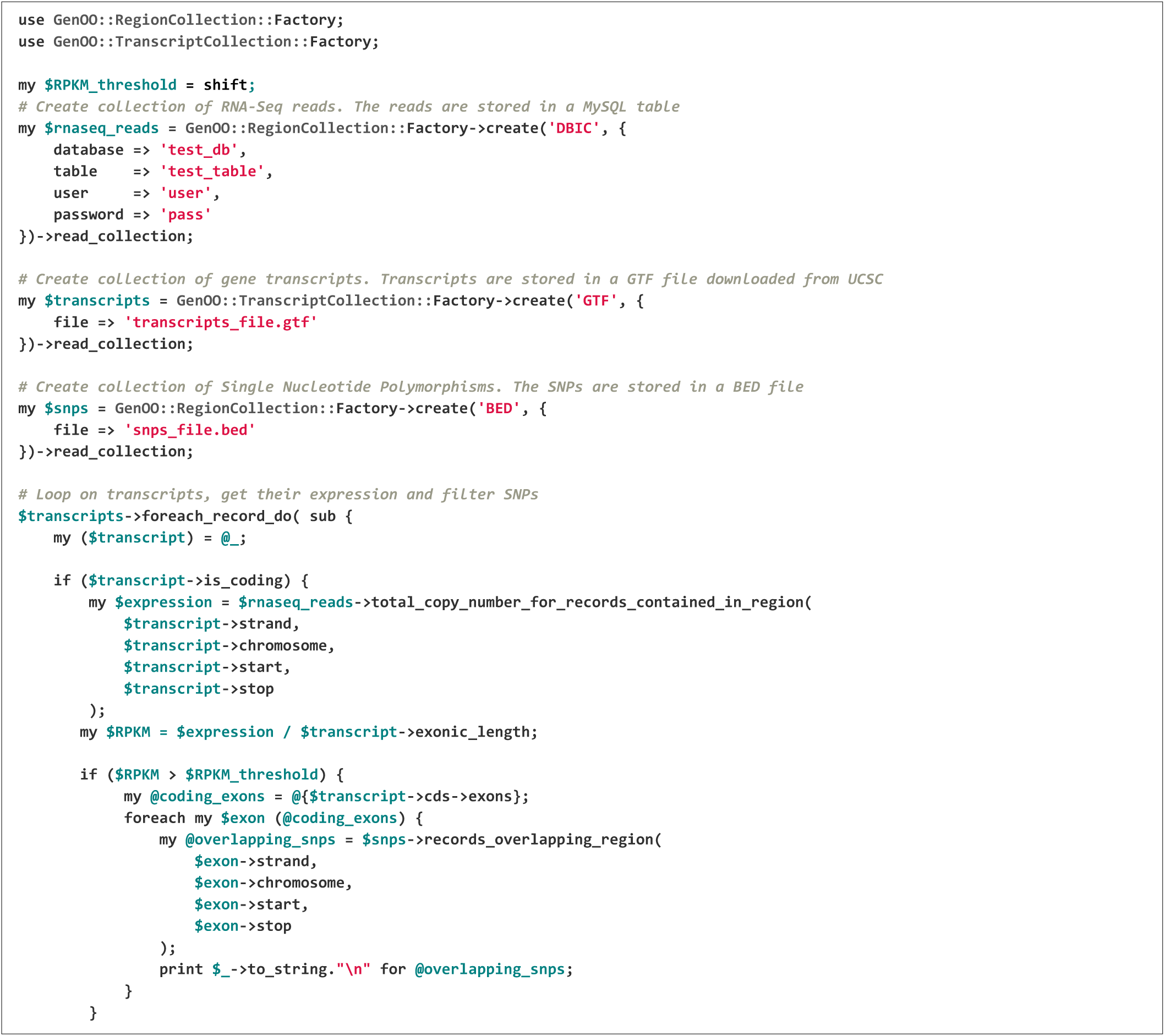
Use case. We filter a list of Single Nucleotide Polymorphisms (SNPs) to keep only those that overlap with the coding sequence of expressed mRNAs as defined by an RNA-Seq experiment

This example illustrates the simplicity with which the four main parts in a typical analysis are performed and how different steps and datasets are integrated together. The first part is the creation of gene models which are read from a GTF file. As shown in the example the gene models offer several convenient functions that enable filtering and extraction of annotation in a natural way. Coding sequences and the location of coding exons are easily retrieved from the gene model.

The second and third part show how the records (reads/SNPs) that are stored in a database or in a BED file are represented as collections in the GenOO framework offering a common interface for advanced range queries and iterations.

The fourth and most important part illustrates how to use a collection and how to find the records of a collection that overlap with a region of interest. This last part is arguably one of the most common tasks performed in a custom HTS analysis and yet still one of the hardest to perform efficiently and correctly. This example illustrates how GenOO provides a common interface to perform this analysis on collections having widely variable underlying structures.

## DISCUSSION

The accelerating rise of HTS as a widely used technique for biological experiments has intensified the need for bioinformatical tools that can efficiently address custom highly specific analyses on HTS data. Some code libraries have been developed that can support the creation of such tools, but their code base is usually fragmented which makes improvements and modifications a daunting affair for developers.

In fact, it is often the case that specific modules within the same code base have been developed under different programming philosophies and conventions, becoming hard to integrate and use efficiently. As an example, BioPerl has a monolithic core and a multitude of classes increasing the complexity of needed modifications and refactoring. Recently, an effort has taken place to break the core of BioPerl into several smaller modules, but the effort is still underway.

Additionally, the scope of these code libraries is to be able to address diverse bioinformatics analysis from proteomics to image manipulation to phylogenetic and genomic analyses. In effect, this creates a large overhead for researchers that specifically need to work on HTS analyses, as they have to deal with often useless and confusing functionality, intended for completely different purposes. Incomplete or outdated documentation exacerbates the problem.

A common reaction to the above mentioned issue is for bioinformatics development groups to rewrite analysis specific code from the beginning, sometimes using small parts of the already developed libraries as black boxes. This approach is time consuming, error prone and unless rigorous testing and validation of the tools written can be performed alongside the development it should preferably be avoided.

In contrast, the aim of GenOO is to make the most common, straightforward tasks in HTS analyses as simple and portable as possible and also to support the rapid development of custom tools. In a HTS analysis custom tools can range from a few filtering scripts or parsers that serve as glue between some larger chunks of analysis or can even be more complex scripts that address specific biological question for which no ready-made tool is available. The goal of GenOO is not to be as feature rich in a wide field of analyses such as all-encompassing libraries (e.g. BioPerl, BioPython, BioRuby), but rather to untangle programmers from the complexity that other generic libraries enforce and allow them to rely on properly developed and tested methods instead of homebrewed scripts.

## CONCLUSIONS

GenOO was developed and has matured through a long process as it was extensively used for the analysis of HTS data sets produced by RNA-Seq and HITS-CLIP experiments [12–14]. Through this maturation it has evolved into a stable and high quality tool. Particular focus has been given to the design of the framework implementing modern Perl concepts and making the code base as easy and extensible as possible. This will enable future development and will allow the scientific community to contribute to the code base in an organized way that will reduce fragmentation and interface inconsistencies. We strongly believe that the use of the framework will make HTS analysis tool programming more efficient and less error prone. It will also help standardize methods, increase their reusability, create code of higher overall quality and help coders avoid common pitfalls and complexities inherent in HTS analyses.

## AVAILABILITY AND REQUIREMENTS

Project name: GenOO

Project home page: https://github.com/genoo/GenOO

Operating system(s): Platform independent

Programming language: Perl

License: Perl

## AUTHORS’ CONTRIBUTIONS

MM designed the framework, developed the framework and drafted the manuscript. PA designed the framework, developed the framework and drafted the manuscript. ZM designed the framework and drafted the manuscript.

## ACKNOWLEDGEMENTS

We would like to thank all the members of the Mourelatos lab for their helpful discussions and comments.

*Funding:* Supported by NIH grants GM072777, NS072561, NS056070 to Z.M. and a Brody Family Medical Trust Fellowship to M.M.

